# Latest RNA and DNA nanopore sequencing allows for rapid avian influenza profiling

**DOI:** 10.1101/2024.02.28.582540

**Authors:** Albert Perlas, Tim Reska, Guillaume Croville, Ferran Tarrés-Freixas, Jean-Luc Guérin, Natàlia Majó, Lara Urban

## Abstract

Avian influenza virus (AIV) currently causes a panzootic with extensive mortality in wild birds, poultry, and wild mammals, thus posing a major threat to global health and underscoring the need for efficient monitoring of its distribution and evolution. Here, we utilized a well-defined AIV strain to systematically investigate AIV characterization through rapid, portable nanopore sequencing by (i) benchmarking the performance of fully portable RNA extraction and viral detection; (ii) comparing the latest DNA and RNA nanopore sequencing approaches for in-depth AIV profiling; and (iii) evaluating the performance of various computational pipelines for viral consensus sequence creation and phylogenetic analysis. Our results show that the latest RNA-specific nanopores can accurately genomically profile AIV from native RNA while additionally detecting RNA epigenetic modifications. We further identified an optimal laboratory and bioinformatic pipeline for reconstructing viral consensus genomes from nanopore sequencing data at various rarefaction thresholds, which we validated by application to real-world environmental samples for AIV monitoring in livestock.

**Author Summary:** We tested portable, rapid, and easy-to-use technology to obtain more information about the potentially zoonotic RNA virus avian influenza virus, or AIV. AIV has spread globally via the migratory paths of wild birds, and endangers domestic birds, mammals, and human populations given past evidence of infections of different animal species. We here used novel genomic technology that is based on nanopores to explore the genomes of the virus; we established optimized ways of creating the viral genome by comparing different laboratory and computational approaches and the performance of nanopores that either sequence the viral RNA directly or the converted DNA. We then applied the optimized protocol to dust samples which were collected from a duck farm in France during an AIV outbreak. We showed that we were able to use the resulting data to reconstruct the relationship between the virus responsible for the outbreak and previously detected AIV. Altogether, we showed how novel easy-to-use genomic technology can support the surveillance of potentially zoonotic pathogens by accurately recreating the viral genomes to better understand evolution and transmission of these pathogens.

## Introduction

Avian influenza virus (AIV) currently causes the largest and deadliest panzootic on the European and American continents [1]; it is known to have spilled over from wild bird populations to poultry and humans, posing a risk for causing a future pandemic [2]. Wild birds are the main reservoir of low-pathogenicity AIV (LPAIV), in particular the Anseriformes and Charadriiformes orders [3]. These birds are asymptomatic to LPAIV and can spread the virus to poultry around the globe [4]. Once in gallinaceous species, LPAIV can evolve into high pathogenicity AIV (HPAIV), resulting in animal welfare, financial and social issues due to high poultry mortality, economic loss, and food insecurity [1]. LPAIV and HPAIV further have the potential to adapt and spread to mammalian species. Since the emergence of H5N1 HPAIV in a domestic goose in Guangdong China in 1996 (“Gs/GD lineage”), it has become clear that HPAIV can also be transmitted back to and subsequently maintained in wild bird populations [5]. As many Anseriformes and Charadriiformes populations perform long-distance migrations, they can rapidly spread AIV variants across countries and continents [6].

AIV is a segmented, negative-strand RNA virus from the *Orthomyxoviridae* family. Its error-prone polymerase, and therefore high mutation rate, as well as its segmented genome in combination with mixed infections allow this virus to be in continuous evolution due to antigenic drift and antigenic shift [4]. One such example is the frequent mutation of LPAIV into HPAIV after recurrent replication in poultry, which provides the perfect environment for the virus to mutate due to the high density of susceptible, genetically similar hosts [7]. This evolutionary plasticity of AIV means that the application of fast genomic technologies to determine their nucleotide composition can help to quickly characterize AIV genomic variation including low-frequency variants, predict virulence, reconstruct transmission dynamics, and determine an outbreak’s origin [8].

The application of *in situ* real-time nanopore sequencing technology by Oxford Nanopore Technologies provides a currently unique genomics-based approach to characterize AIV in a fast, straightforward, and cost-efficient manner all around the world [9], which makes viral surveillance accessible in low- and middle-income countries as well as in remote field setting for wild bird monitoring. This technology has been established for AIV profiling through sequencing of complementary DNA (cDNA) after retro-transcription (RT) and multi-segment PCR amplification (M-RTPCR) [8,10,11]. While ligation-based and transposase-based rapid sequencing library preparations (“DNA-nanopore” chemistry R9) have been applied to nanopore-sequence cDNA from AIV [12], a variety of computational pipelines have subsequently been used for data analysis and consensus sequence generation, which have however not yet been systematically assessed and compared [8,10,13–17]. Keller et al. [14] have further applied direct RNA nanopore sequencing to the viral RNA (vRNA) of AIV, which could circumvent biases introduced through cDNA synthesis [18]. This protocol is faster due to the omission of M-RTPCR, and further allows for the detection of RNA modifications [19]. Direct RNA sequencing through nanopore technology (DNA-nanopore chemistry R9) has, however, suffered from high sequencing error rates as well from high RNA input requirements and from a lack of multiplexing options for efficient sample processing [14,20].

Here, we used a well-defined viral culture to conduct a systematic study for AIV characterization through nanopore sequencing by (i) comparing cDNA and vRNA sequencing of AIV in terms of sequencing data throughput, data quality and consensus sequence accuracy, and by (ii) systematically assessing the performance of different computational analysis pipelines. Besides the current gold standard nanopores for DNA (DNA-nanopore chemistry R10) and RNA (DNA-nanopore chemistry R9) sequencing, we have for the first time applied RNA-specific nanopores (“RNA-nanopore” chemistry) for RNA virus profiling; this chemistry is based on completely new nanopores that have been optimized for direct RNA sequencing – in contrast to the previous DNA-nanopore chemistry R9 which relies on nanopores optimized for DNA sequencing and therefore suffers from a high sequencing error rate of >10% [14]. We further included several protocols to compare portable approaches for *in situ* applications with standard laboratory-based approaches. Finally, in order to test the application of nanopore sequencing to samples with lower AIV loads such as environmental samples, we optimized our approaches through data rarefaction simulations, and finally used our results to characterize AIV from non-invasively collected dust samples.

## Results

### Similar performance of laboratory-based and portable RNA extraction and quantification protocols

Using LPAIV H1N1 viral cultures, we found that the NucleoSpin RNA Virus kit was the most efficient RNA extraction approach, yielding the lowest Ct (cycle threshold) values. While we therefore continued our analyses with this kit, the portable Biomeme M1 Sample Prep Cartridge Kit, which allows for RNA extraction in just 5 minutes, yielded only slightly higher Ct values (S1). The standard RT-PCR and portable Mic qPCR systems further showed comparable performance (S1); we therefore continued our analyses with the Mic qPCR machine.

### Nanopore sequencing genome coverage and rarefaction analysis

While the sequencing read length distributions were largely consistent across the cDNA, RNA002, and RNA004 nanopore sequencing approaches (Methods), the RNA002 dataset comprised a lower number of sequencing reads (**Fig 1A**). The alignment of the reads to the AIV reference segments further showed an uneven coverage distribution across the genome, with all sequencing approaches leading to similar coverage of the polymerase segments; every sequencing approach further led to increased coverage at the ends of each segment (**Fig 1B**).

**Fig 1.**
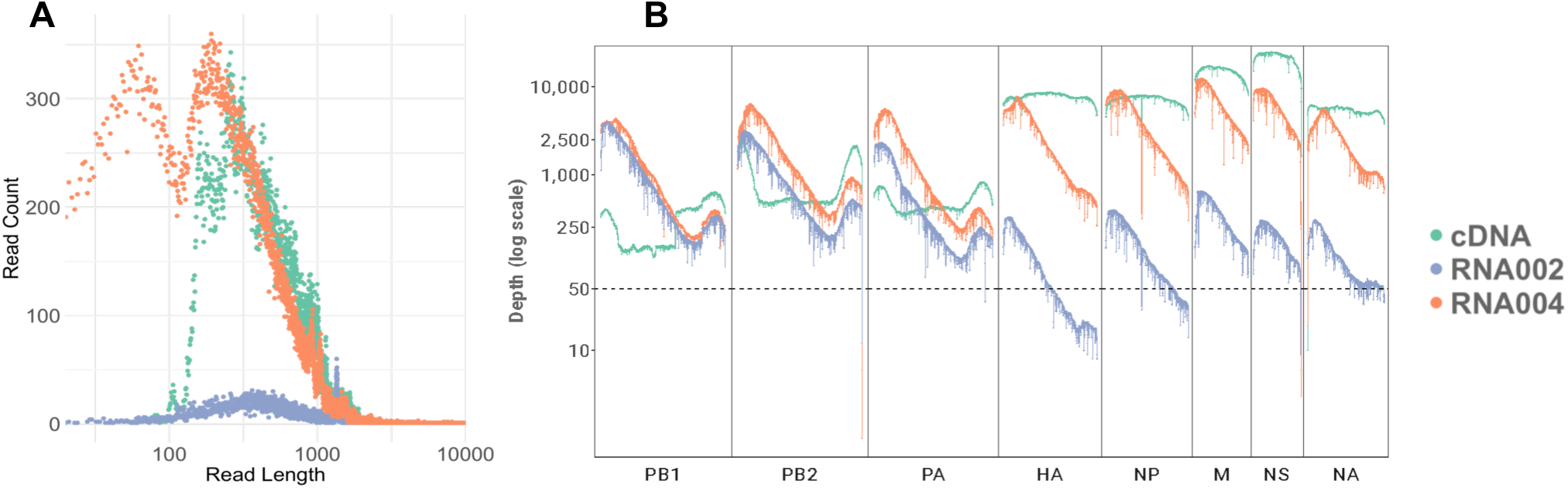
Nanopore sequencing results of an AIV viral culture using DNA-nanopores (“cDNA” sequencing through R10 chemistry; direct RNA sequencing through “RNA002” R9 chemistry) and RNA-nanopores (direct RNA sequencing through “RNA004” RNA chemistry). **A.** Sequencing read length distribution across the cDNA, RNA002, and RNA004 datasets. **B.** Reference genome coverage of the three sequencing datasets across all AIV segments (PB1: Polymerase basic 1, PB2: Polymerase basic 2, PA: Polymerase acidic, HA: Hemagglutinin, NP: Nucleoprotein, NA: Neuraminidase, M: Matrix, NS: Nonstructural). The horizontal line indicates a coverage of 50x.

Given the uneven throughput and coverage across the three sequencing datasets, we performed rarefaction for all downstream analyses and re-assessed the AIV genome coverage (Methods; **Table 1**; **S2**).

**Table 1.**
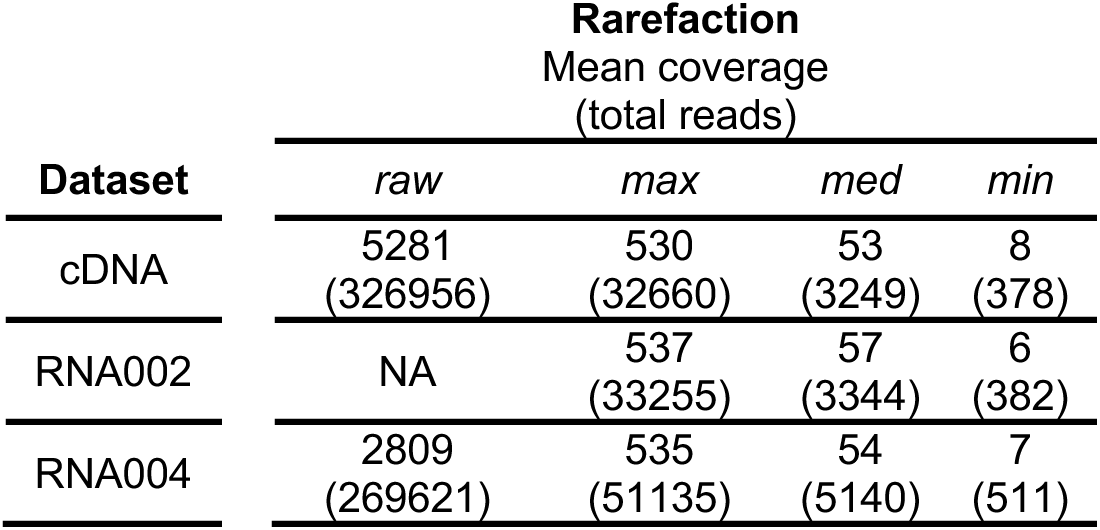
Mean AIV reference genome coverage and (in brackets) total number of reads across each sequencing dataset (rows: cDNA, RNA002, RNA004) and respective rarefaction (columns: *raw* for total dataset; *max* for same mean coverage; *med* for 10% of the *max* data; *min* for 1% of the *max* data).

### AIV consensus sequence creation

We first used the BIT score to evaluate the quality of the viral consensus sequence created by different computational pipelines in comparison to the known AIV reference (Methods). Even at maximum coverage (*max* rarefaction), only the reference-based approaches BCFtools or iVar, and the iterative reference-based assembly tool IRMA were able to create the full consensus sequence of all eight viral segments (**Fig 2A**). Reference-based EPI2ME did not assemble the NS and HA segments. The hybrid approach CZID as well as the *de novo* assembler metaFlye only assembled the largest segments PA, PB1, and PB2, and did not create consensus sequences of the unassembled reads mapping to the other segments. Flye (without the metagenomics configuration) only assembled the PA and PB2 segments.

**Fig 2.**
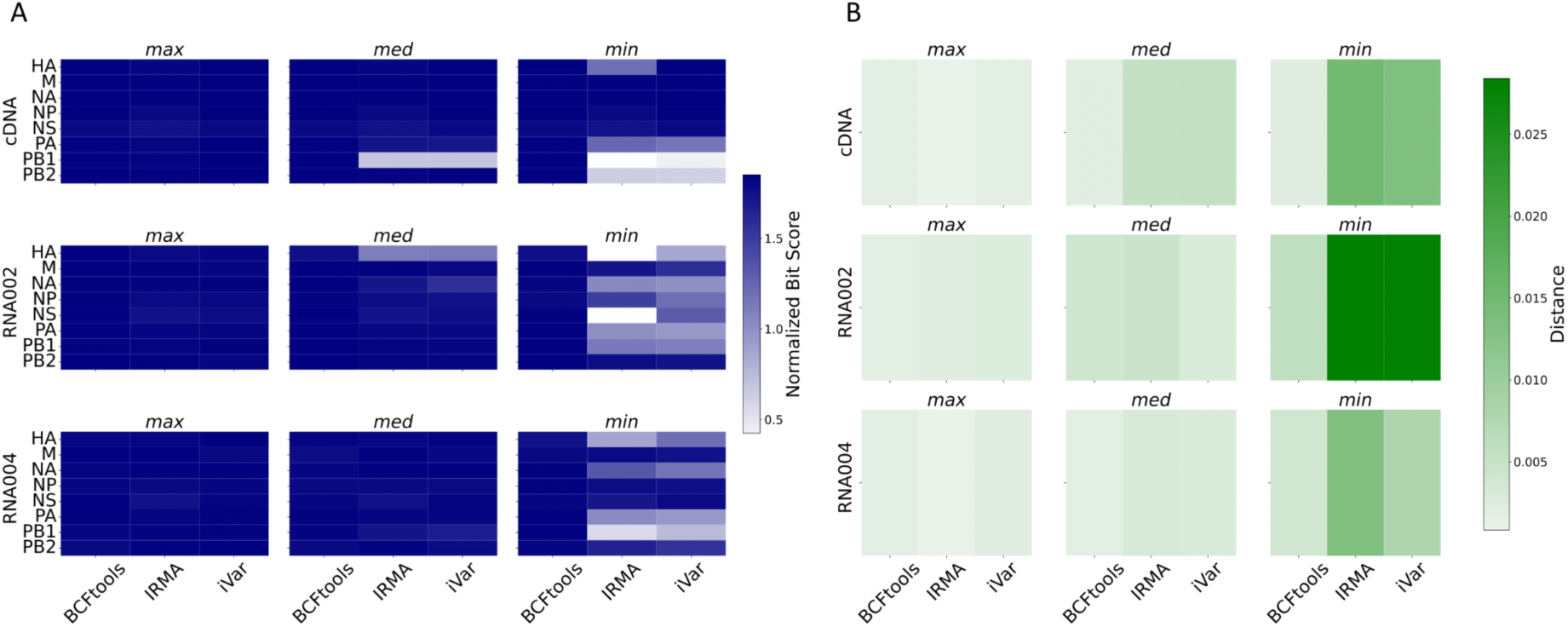
Evaluation of viral consensus sequence creation from nanopore sequencing datasets (cDNA, RNA002, RNA004) across all data rarefactions (*min*, *med*, *max*). The performance of the computational tools BCFtools, iVar, and IRMA, which were the best-performing approaches for the *max* datasets, is visualized. **A.** Consensus sequence evaluation across the eight viral AIV segments through normalized BIT scores calculated based on the known AIV reference. **B.** Consensus sequence evaluation through whole-genome evolutionary distance comparisons with the known AIV reference.

For our rarefied datasets (*med* and *min*), we found further performance differences between BCFtools, iVar, and IRMA (**Fig 2A**). For all three sequencing approaches (cDNA, RNA002, RNA004), BCFtools performed best across all viral segments, with IRMA being unable to create certain segments at all at *min* rarefaction, namely PB1 for cDNA, and HA and NS for RNA002. Across the sequencing approaches, we only found differences at *min* rarefaction where RNA004 outperformed RNA002, and cDNA surpassed both RNA-based methods with the exception of the polymerase segments.

We next calculated the evolutionary distance between the whole-genome consensus sequences and the known AIV reference (Methods). We found that BCFtools again performed best in that it achieved the smallest evolutionary distance from the true reference across all nanopore sequencing approaches and rarefaction thresholds (**Fig 2B**). The good performance of BCFtools was especially pronounced at *min* rarefaction, where it clearly outperformed the other consensus sequence creation tools. Across the nanopore sequencing approaches, the RNA-nanopore-based RNA004 and cDNA-based approaches worked equally and were quite robust to the viral coverage; the worse performance of the DNA-nanopore-based RNA002 approach was most noticeable for the minimum-coverage data (*min* rarefaction) (**Fig 2B**).

### Nanopore sequencing-based AIV profiling in environmental samples

Given the good performance of cDNA and RN004 for viral consensus sequence creation also from smaller genomic datasets (*min* rarefaction), we next simulated sequencing data from environmental samples by rarefaction to 0.01% (*env_max*) and 0,001% (*env_min*). We focused this analysis on HA as the viral segment that is normally used for AIV lineage identification via phylogenetic analysis, and which plays an important role in host cell penetration. We found that the cDNA sequencing approach was very robust to extremely low viral coverage, yielding lower evolutionary distances of the HA segment to its known reference. Specifically, with *env_max* at a distance of 0.0000025 and a mean coverage of 76, and *env_min* at a distance of 0.00000025 and a mean coverage of 10, these outcomes were significantly better than those achieved through direct RNA sequencing using RNA004. For RNA004, *env_max* resulted in a distance of 0.0012 with a mean coverage of 24, and *env_min* resulted in a distance of 0.0058 with a mean coverage of 5.

Given the good performance of AIV cDNA sequencing and BCFtools analysis for simulated environmental samples, we finally used this approach to process real environmental samples, namely four dust samples from a turkey farm in France. The samples ranged from Ct values of 24 to 26, and all resulted in similar read length distributions and coverage distributions across viral segments (S3). BCFtools was able to create consensus sequences of all eight viral segments, and an evolutionary based on known AIV strains from the NCBI influenza database (Methods) elucidated the phylogenetic relationship between the farm’s viral strain and known AIVs (**Fig 3**).

**Fig 3.**
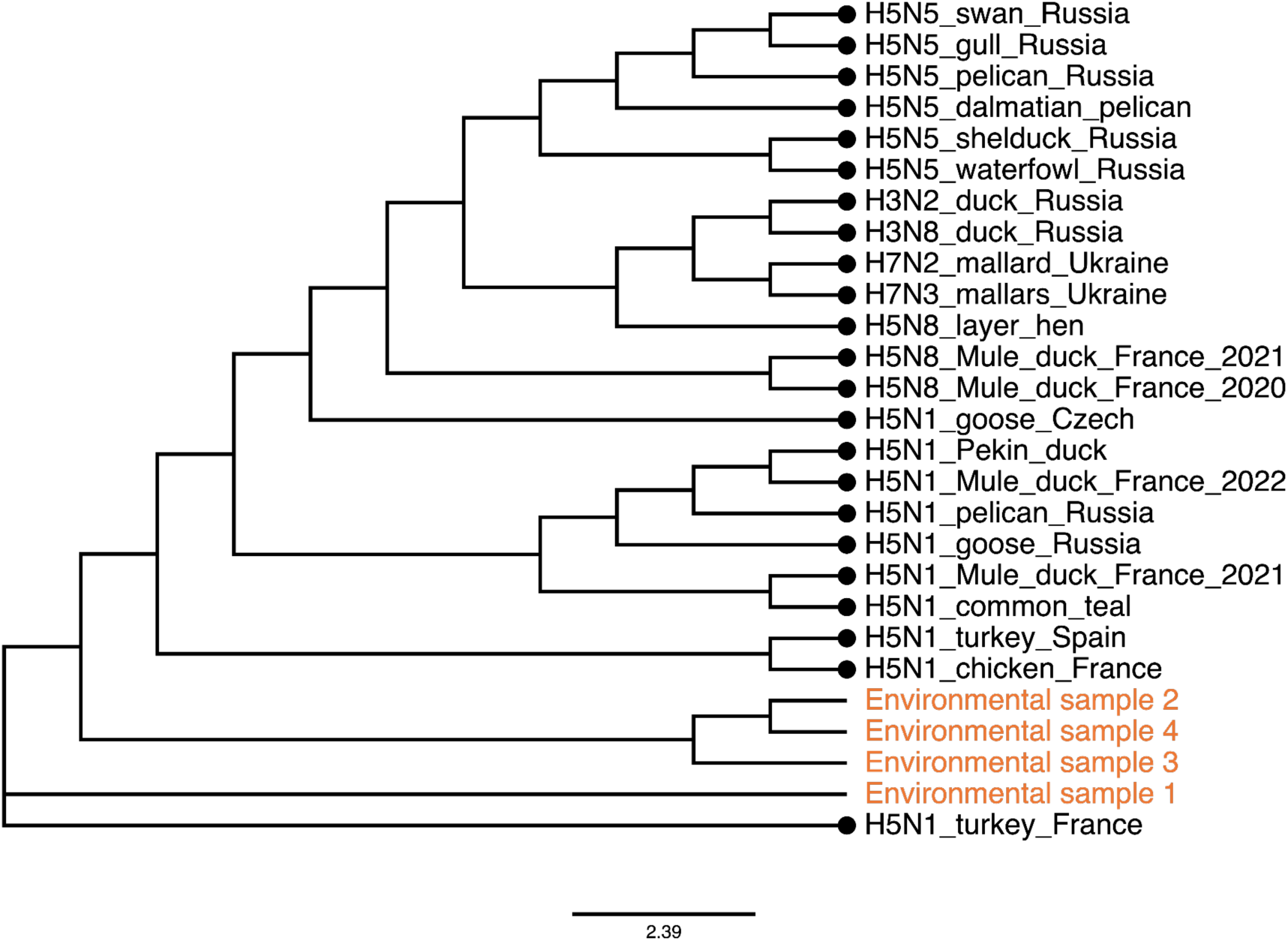
Phylogenetic tree of AIV consensus HA segments from four environmental samples (dust samples from turkey farm in France) and known European AIV strains. AIV of the environmental samples was assessed by cDNA nanopore sequencing, and consensus sequence was created through BCFtools.

### RNA modification profiling

Finally, we identified m6A modifications in the raw RNA004 nanopore data as the best-performing direct RNA sequencing approach in our study. In total, we identified 2145 modifications, with the distribution of total modifications and modifications per base across segments as follows: HA (311 modifications, 0.179 modifications per base), M (227 modifications, 0.230 modifications per base), NA (307 modifications, 0.216 modifications per base), NP (253 modifications, 0.165 modifications per base), NS (185 modifications, 0.208 modifications per base), PA (237 modifications, 0.108 modifications per base), PB1 (282 modifications, 0.122 modifications per base), and PB2 (343 modifications, 0.149 modifications per base). The M segment exhibited the highest modification density, while the PA segment displayed the lowest.

## Discussion

Here, we present an optimized nanopore sequencing pipeline suitable for rapid field studies from non-invasively collected environmental samples. Our protocols aim to identify AIV strains as well as their evolution processes and potential transmission patterns. The implementation of such strategy for AIV monitoring – including in remote areas along long-distance migration routes of potential avian hosts – is very promising for rapidly and appropriately informing control measures as part of a “One Health” strategy.

Our study shows that direct viral RNA as well as cDNA nanopore sequencing provide robust genomic approaches to rapidly create viral consensus sequences *in situ*, to assess the virus’ evolutionary trajectory. While previous studies have explored the application of nanopore sequencing to AIV, they have often been limited to one or a few processing pipelines [8,10,13–17]. Here we identify the optimal computational analysis pipeline for robust analysis of viral data across a range of simulated viral loads.

While direct RNA sequencing has previously been applied to AIV analysis [14], the high sequencing error rate of these protocols, that are based on standard DNA-nanopores, impeded meaningful analyses of the data. Here we find that the latest direct RNA nanopore sequencing technology (which is based on a unique RNA-nanopore specifically designed for transcriptomic rather than genomic research), provides similar results to cDNA sequencing using Oxford Nanopore Technologies’ established high-accuracy DNA-nanopores (R10 chemistry). Our study is therefore, to the best of our knowledge, the first to show that viral genomes can be profiled with high accuracy directly from their RNA without any cDNA reverse-transcription which is laborious, time-consuming and might introduce biases. We also show that direct RNA sequencing simultaneously allows for RNA modification calling [19]. To the best of our knowledge, this study marks the first in-stance of identifying m6A modifications in AIV using direct RNA nanopore sequencing technology. Such modifications play a critical role in viral RNA viruses, allowing them to mimic host RNA and thereby evade the host’s immune system, which underscores the significance of direct RNA sequencing for epidemiology and immunology [21,22].

We compared the performance of several reference-based, *de novo* assembly, and hybrid computational approaches to reconstruct the viral consensus sequence from nanopore data at various rarefaction thresholds. While web-based tools such as EPI2ME and CZID are more user-friendly than the remaining tools which rely on the usage of the command line, they did not perform well in creating the consensus sequence of all AIV segments – even in the datasets with a high genome-wide coverage of >500x. In the case of the reference-based EPI2ME tool, the poor performance could be due to the analysis’ restriction to US AIV references. In the case of the hybrid-assembler CZID, only assemblies from the longer viral segments could be obtained while many un-assembled reads aligned to the reference sequences of the smaller segments. We faced the same problem when using the *de novo* assembly command line tool Flye, which can be explained by Flye’s incompatibility with reads shorther than 1 kb, which exceeds the entire length of some viral segments. We found that the Flye version for metagenome assembly (using the –meta flag) worked better for reference reconstruction; this might be related to the fact that metaFlye does not assume even coverage across the genome, which is a suitable configuration for highly diverse RNA viruses where amplification or targeting biases might result in uneven coverage across segments [23–25].

The reference-based command line tools BCFtools and iVar as well as IRMA, which relies on iterative refinement, were able to create high-quality viral consensus sequences for all nanopore data if available at high-coverage. Although we could have evaluated other reference-based tools, we chose to utilize the most commonly employed ones for generating consensus sequences from AIV in this study. While some of these computational pipelines have previously been applied to AIV nanopore sequencing data, they have not yet been compared to each other, especially in application to different nanopore sequencing modalities [13,14,16,17]. Rarefaction of this high-coverage viral data identified BCFtools as the best tool in generating consensus sequences across viral segments similar to the known reference (measured by high BIT scores and small evolutionary distances). The good performance of BCFtools throughout our analyses might be due to its – in comparison to iVar and IRMA – relatively strong reliance on reference data [26,27] and our incorporation of a comprehensive reference database. This means that the performance of BCFtools might worsen in the case of highly divergent and previously unseen RNA viruses.

Given the good performance of BCFtools in combination with cDNA and RNA004 nanopore sequencing data, we further simulated AIV sequencing from environmental samples. We show that cDNA sequencing leads to relatively higher BIT scores and smaller evolutionary distances than RNA004 sequencing for such low-concentration samples, which might be due to the still relatively decreased sequencing accuracy of direct RNA sequencing in comparison to direct DNA sequencing (96% for RNA004 using the RNA chemistry, vs. 99% for cDNA using the R10 chemistry;). When testing our AIV profiling pipeline on real environmental samples, namely dust samples from duck farms, we therefore employed the combination of cDNA sequencing and BCFtools-based analysis. We were able to reconstruct complete viral consensus sequences from this data, which we leveraged to reconstruct a phylogenetic tree and to show evolutionary similarity between our AIV strains and contemporary H5 AIV strains. These H5 strains are responsible for the ongoing severe HPAIV panzootic [1], and one of the most closely related strains has actually been responsible for a H5 HPAIV outbreak in another french farm.

An additional challenge for the analysis of low-concentration viral samples can be the uneven segment coverage that we observed across and within viral segments. Our cDNA data showed decreased coverage of the polymerase segments, while the RNA002 data showed decreased coverage of the respective other segments. We hypothesize that these coverage disparities stem from biases introduced through the use of universal primers for cDNA amplification and through the oligo-nucleotide adapters targeting AIV for direct RNA sequencing, respectively. The newest direct RNA sequencing protocol RNA004, on the other hand, relies on an alternative ligase enzyme, which might explain its more even coverage across segments. Within segments, all nanopore sequencing approaches result in uneven coverage. This especially applies to the direct RNA sequencing approaches, where the systematic decrease in coverage towards the end of the segment might be explained by sequencing adapters targeting the segments’ conserved 3’-end and by rapid RNA fragmentation [14].

Our study additionally showcases the field applicability of our nanopore sequencing protocols by benchmarking fully portable equipment. While we found that standard column-based viral RNA extraction outperformed more portable alternatives, the Biomeme M1 Sample Prep Cartridge approach only led to slightly increased Ct values, suggesting its potential for future field studies. We further show that the MIC qPCR [8], MinION MK1c nanopore sequencing device, and rapid library preparation protocols provide a fully portable framework to conduct AIV profiling at the point of interest all around the world, even without internet access.

## Methods

### RNA extraction and quantification

H1N1 LPAIV was isolated from a duck sample in 2006 (strain A/duck/Italy/281904/2006) and isolated in specific pathogen-free (SPF) eggs as previously described [30]. The high-quality reference genome was obtained from Sanger sequencing data ([31], GenBank accession number: FJ432771). We extracted RNA from egg allantoic fluids using Macherey-Nagel’s NucleoSpin RNA Virus extraction kit, and quantified the extracted RNA using the Qubit RNA BR assay. We additionally used Biomeme’s M1 Sample Prep Cartridge Kit For RNA 2.0 [8] and Lucigen’s Quick Extract DNA Extraction Solution kit to assess the performances of faster and portable RNA extraction approaches. For Quick Extract, we followed the manufacturer’s instructions and, additionally, an alternative method adapted for SARS-CoV-2 RNA extraction [32]. We then compared the performance of the different kits in terms of detection and quantification rates using standard RT-PCR (Applied Biosystems 7500 Fast Instrument, Thermo Fisher) and portable RT-PCR (Magnetic Induction Cycler quantitative PCR (Mic qPCR), Bio Molecular Systems). We targeted a highly conserved region of 99 bases of the AIV MP gene using previously established approaches to detect and quantify AIV using RT-PCR [33,34].

### Nanopore sequencing

We then performed nanopore sequencing of the NucleoSpin RNA extracts. First, we performed direct vRNA sequencing using the DNA-nanopore (R9 chemistry; “RNA002” kit) and the RNA-nanopore chemistry (RNA chemistry; “RNA004” kit). We specifically targeted AIV RNA following the protocol described by Keller et al. [14]. Briefly, direct RNA nanopore sequencing requires a reverse transcriptase adapter (RTA) which usually captures poly(A) tails of the messenger RNA (mRNA); a sequencing adapter then ligates to the RTA and directs the mRNA to the nanopore. To target AIV RNA, we used a modified RTA, i.e. a custom oligo-nucleotide that is complementary to the 3’-region that is conserved across all AIV segments. As these conserved regions differ slightly across segments, we used two custom oligo-nucleotides, RTA-U12 and RTA-U12.4, which were mixed at a molar ratio of 2:3 to a total concentration of 1.4μM [14]. We subsequently used the portable MinION Mk1c device for nanopore sequencing; for the R9 chemistry sequencing, we used a FLO-MIN106 R9.4.1 flow cell, and for the RNA chemistry, we used a FLO-MIN004RA flow cell.

Second, we performed cDNA sequencing using the latest DNA-nanopore chemistry (R10 chemistry) and rapid barcoding library preparation (SQK-RBK114.24) after cDNA conversion of the extracted RNA and multi-segment amplification through M-RTPCR. M-RTPCR was performed as described previously, targeting the conserved regions across all AIV segments [35,36]. Briefly, the extracted RNA was mixed with Superscript III One-Step PCR reaction buffer and the previously defined primers, the PCR reactions were run on a portable Mic qPCR device. For sequencing, we used three barcodes with the same sample to increase the total quantity of cDNA added to the final sequencing library. We subsequently used the portable MinION Mk1c device and a FLO-MIN114 R10.4.1 flow cell for nanopore sequencing.

### Data processing

We obtained raw nanopore sequencing data in fast5 format for the DNA-nanopore, and in pod5 format for the RNA-nanopore sequencing runs. For the DNA-nanopore runs, we used the Guppy (v6.4.8+31becc9) high-accuracy basecalling model (HAC; rna_r9.4.1_70bps_hac model for vRNA, dna_r10.4.1_e8.2_400bps_hac model for cDNA); for the RNA-nanopore run, we used the Dorado (v0.4.3+656766b) HAC model for RNA (rna004_130bps_hac). After removing short reads (<50 bases) using SeqKit (v2.4.0) [37], we used Minimap2 (v2.26) [38] with the *-ax map-ont* configuration for cDNA and the *-ax splice -uf -k7* configuration for vRNA reads to align the resulting fastq files to our ground-truth reference genome (GenBank accession number: FJ43277). We converted the resulting sam files to bam files, indexed, and sorted them using SAMtools (v1.17) [39] to obtain the genome coverage distribution.

### Data rarefaction

To compare all nanopore sequencing results, we rarefied the three genomic datasets (cDNA, vRNA by DNA-nanopore: RNA002, vRNA by RNA-nanopore: RNA004) from the “*raw*” data to the same mean coverage (“*max*” data). After rarefying the cDNA fastq file to 10% and the RNA004 fastq file to 20% of its original number of reads, a similar mean coverage to the RNA002 fastq file was achieved (mean genome coverage of 537). We further rarefied this *max* data to simulate results from samples with lower viral load, namely to 10% of the *max* data (“*med*”) and to 1% of the *max* data (“*min*”). Finally, to simulate real environmental samples with potentially extremely low viral loads, we additionally rarefied the raw cDNA and RNA004 data to 0.01% (“*env_max*”) and 0.001% (“*env_min*”) for follow-up analyses.

### Consensus sequence construction

For reference-based consensus sequence creation, we mapped each dataset to a reference database generated for each segment from the NCBI Influenza Virus Database, which contains all AIV nucleotide sequences from Europe (as of 04/03/2023). We excluded the true reference sequence of our H1N1 virus from all segment-specific reference databases in order to simulate a realistic situation where the true genomic sequence of our AIV strain would not yet be known. We indexed the reference databases and mapped our sequencing reads against the databases using Minimap2. We then indexed and sorted the sam files and converted them to bam files using samtools. Using samtools idxstats, we selected the reference to which most reads mapped across segments. All our reads were then mapped to the full best reference genome using Minimap2. We then tested two standard reference-based computational pipelines to create the consensus sequence from this alignment, BCFtools (v1.17) [26] and iVar (v1.4.2) [27].

We additionally used the Iterative Refinement Meta-Assembler (IRMA; v1.0.3) [40] that iteratively refines the reference used in the analysis to increase the accuracy of the consensus sequence obtained. Using this pipeline, the consensus of each segment can be obtained directly from the fastq file without intermediate steps required by the user. We used the “FLU-minion” configuration for nanopore sequencing data, which drops the median read Q-score filter from 30 to 0, raises the minimum read length from 125 to 150, raises the frequency threshold for insertion and deletion refinement from 0.25 to 0.75 and 0.6 to 0.75, respectively, and lowers the Smith-Waterman mismatch penalty from 5 to 3 as well as the gap open penalty from 10 to 6.

We further applied Oxford Nanopore Technologies’ EPI2ME (v.5.1.9.) workflow for influenza viruses (“wf-flu”) to our data, which is also based on a reference-based consensus sequence creation approach, but which uses a specific influenza reference database which only focuses on the FluA and FluB segments (https://labs.EPI2ME.io/influenza-workflow/).

For *de novo* consensus sequence creation, we used Flye (v2.9.2) with and without – meta flag [41] followed by assembly polishing using racon (v1.4.3) [42]. We additionally applied the Chan Zuckerberg ID (CZID) [43,44] pipeline to our data, which performs a combination of *de novo* and reference-based approaches: It uses metaFlye to assemble the data and generate contigs, followed by Minimap2-alignments of the still unassembled reads against the NCBI database [45].

To evaluate the consensus sequence creation pipelines, we first used blastn (v2.15) [46] to align every consensus segment to our known reference and we normalized the resulting BIT score by segment size to facilitate comparisons across different segments. Then, we assembled the whole-genome consensus sequences by concatenating the segment-specific consensuses derived from each of the rarefied datasets. We next performed a phylogenetic analysis of all reconstructed AIV whole-genome consensus sequences together with the known reference using a maximum likelihood approach as implemented in IQ-TREE (Jukes-Cantor nucleotide substitution model) (v2.0.6) [47] to obtain pairwise likelihood distances between our consensus sequences and the reference.

### Environmental samples

We finally obtained real environmental samples (surface dust collected with dry wipes on building’s walls and feeders) from 4 HPAIV H5N1 Gs/GD lineage outbreaks in 2022 and 2023 in duck farms in South-west and West regions of France [48]. The environmental samples were processed and analyzed as described above for the LPAIV H1N1 viral cultures. For the phylogenetic tree reconstruction, we incorporated all recent AIV strains from Europe (from January 1^st^ 2020 until May 1^st^ 2023) from the NCBI Influenza Virus Database; visualization was done using IROKI [49]. Due to the relevance of the HA segment for host cell penetration and phylogenetic analysis, we exclusively focused this analysis on this segment.

### Detection of RNA modifications

We additionally searched for N^6^-methyladenosine (m6A) RNA modifications in the RNA-nanopore data using the respective Dorado basecalling model (rna004_130bps_sup v3.0.1_m6A_DRACH@v1). Subsequent analysis was performed using Modkit (v0.2.4.)(https://github.com/nanoporetech/modkit).

## Data access

Original fastq files from all the sequencing runs are available on the European Nucleotide Archive (ENA) under the accession number PRJEB72673.

All our computational scripts are available via the GitHub repository *real-time_surveillance_of_avian-influenza*: https://github.com/Albertperlas/Latest-RNA-and-DNA-nanopores-allow-for-rapid-avian-influenza-profiling

## Acknowledgement

We would like to acknowledge Dr. Ana Moreno from the Istituto Zooprofilattico Sperimentale della Lombardia e dell’ Emilia Romagna (Brescia, Italy) who kindly provided us with the A/duck/Italy/281904/2006 LPAIV H1N1 isolate.

## Author contributions

AP conducted the nanopore sequencing experiments and subsequent bioinformatic analysis. TR, GC, FT, and LU provided essential advice on experimental design and bioinformatics. NM and JG collected the samples for the study. AP and LU significantly contributed to the discussion and manuscript writing. All authors, AP, TR, GC, FT, NM, JG, and LU, contributed to writing, reviewing, editing, and approving the final manuscript.

## Supporting Information

**S1 Figure.**
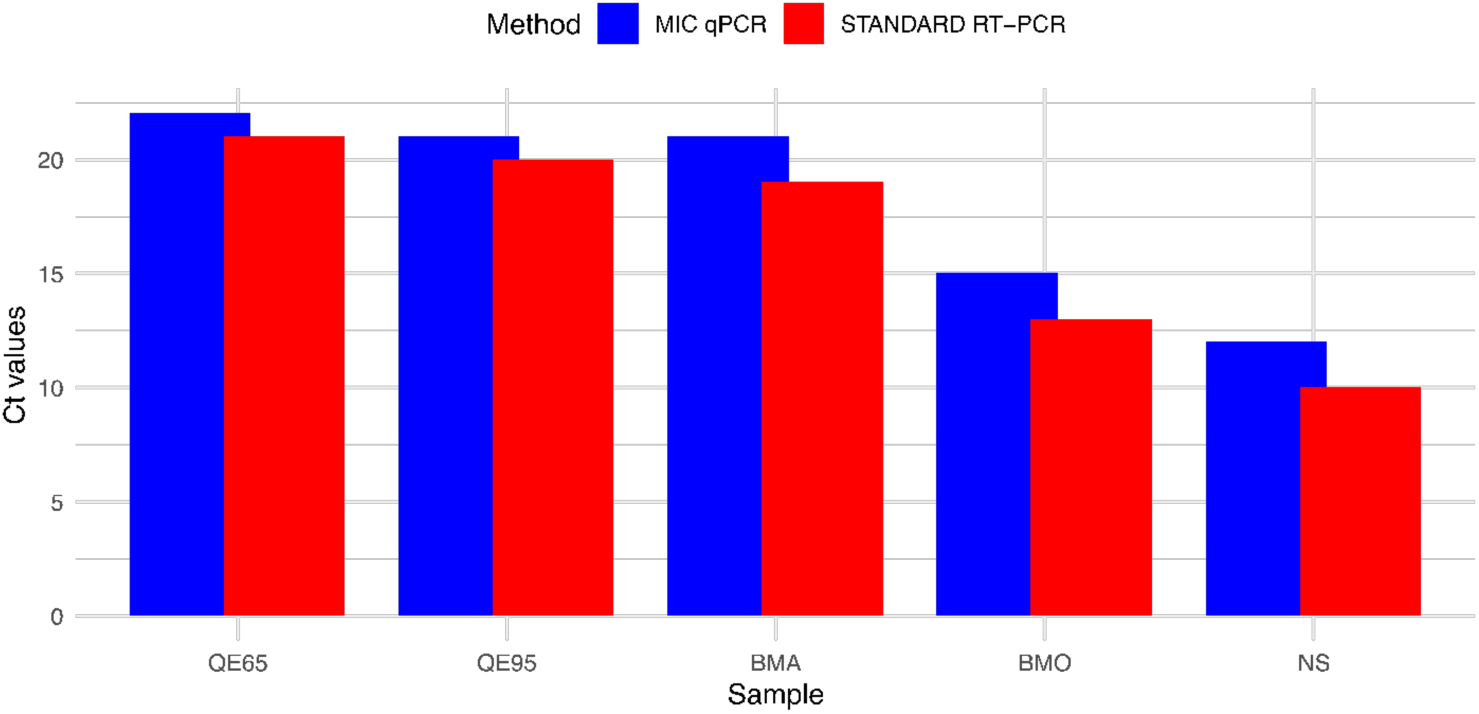
Comparison of Ct values from different RNA extraction kits and quantification methods. The Ct values were determined using the NucleoSpin RNA Virus extraction kit (NS), the Biomeme M1 Sample Prep Cartridge Kit for RNA 2.0 with the manufacturer’s protocol (BMO) and a modified protocol by de Vries et al. (2022) (BMA), and the Quick Extract DNA Extraction Solution with the manufacturer’s protocol (QE95) and an alternative method for SARS-CoV-2 RNA extraction by Ladha et al. (2020) (QE65). Quantification was performed using standard real-time PCR (red columns) and a portable real-time PCR (Mic qPCR, blue columns) targeting the M segment of the virus.

**S2 Figure.**
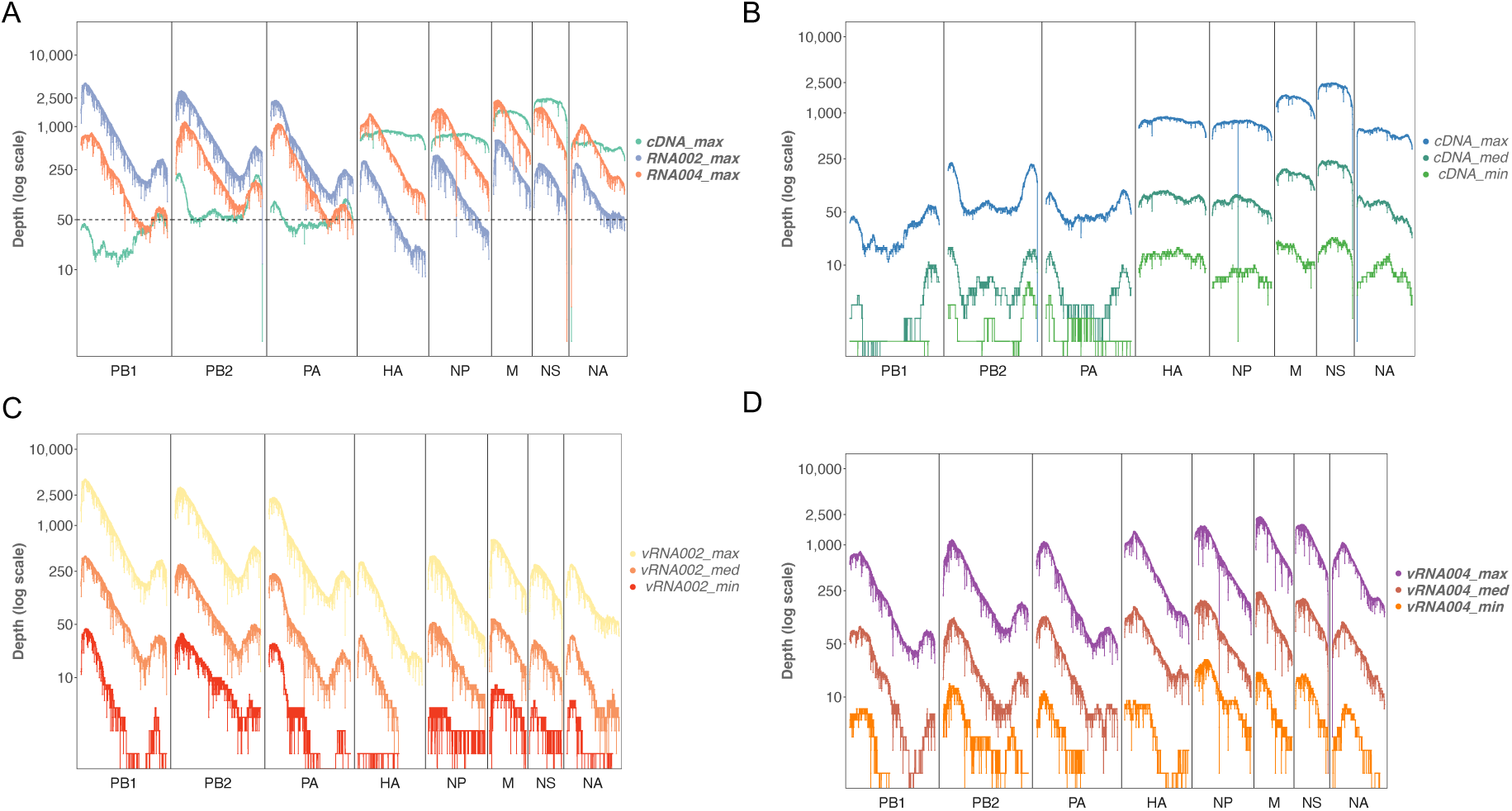
Coverage of the different datasets after rarefaction. **A.** Maximum coverage with similar mean coverage of all datasets (*cDNA-max, vRNA004-max*, and *vRNA002-max*). **B.** Coverage from cDNA datasets after rarefaction (*cDNA-max*, *cDNA-med*, and *cDNA-min*). **C.** Coverage from vRNA002 dataset (*vRNA002-max*) after rarefaction (*vRNA002-med*, and *vRNA002-min*). **D.** Coverage from vRNA004 dataset after rarefaction (*vRNA004-max*, v*RNA004-med*, and *vRNA004-min*).

**S3 Figure.**
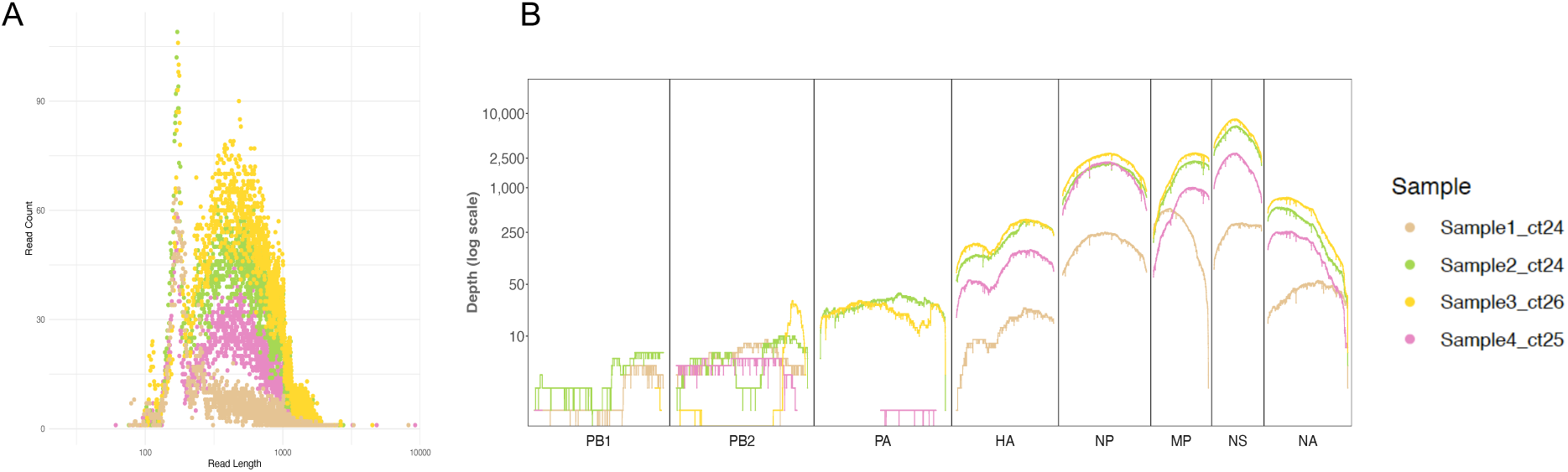
**A.** Read length distribution plots of the environmental samples. **B.** Coverage of each nucleotide position in each segment from cDNA sequencing of our environmental samples.

